# A Mixed-Modeling Framework for Whole-Brain Dynamic Network Analysis

**DOI:** 10.1101/2021.02.25.432947

**Authors:** Mohsen Bahrami, Paul J. Laurienti, Heather M. Shappell, Sean L. Simpson

**Author notes:** **Corresponding Author.** Department of Radiology, Wake Forest School of Medicine, Winston-Salem, NC 27157, USA.

## Abstract

The emerging area of dynamic brain network analysis has gained considerable attraction in recent years. While current tools have proven useful in providing insight into dynamic patterns of brain networks, development of multivariate statistical frameworks that allow for examining the associations between phenotypic traits and dynamic patterns of system-level properties of the brain, and drawing statistical inference about such associations, has largely lagged behind. To address this need we developed a mixed-modeling framework that allows for assessing the relationship between any desired phenotype and dynamic patterns of whole-brain connectivity and topology. Unlike current tools which largely use data-driven methods, our model-based method enables aligning neuroscientific hypotheses with the analytic approach. We demonstrate the utility of this model in identifying the relationship between fluid intelligence and dynamic brain networks using resting-state fMRI (rfMRI) data from 200 subjects in the Human Connectome Project (HCP) study. To our knowledge, this approach provides the first model-based statistical method for examining dynamic patterns of system-level properties of the brain and their relationships to phenotypic traits.

## 1. Introduction

The past two decades have witnessed an explosion of studies aimed at examining the brain as a complex system through analysis of neuroimaging data, particularly data from functional MRI (fMRI). Complex functional systems of the brain are often analyzed through graph theoretical measures of the brain’s functional networks (Bullmore and Sporns 2009). Nodes in brain networks often represent brain regions, and edges represent functional connections (statistical associations) between the blood-oxygen-level-dependent (BOLD) signals in different brain regions. Until recent years, most network studies of the brain focused on static functional networks, in which the functional connections between brain regions were defined over the entire scanning period or condition. Although such studies have provided promising insights into functional organization and abnormalities of the brain (Bassett and Bullmore 2009, Park and Friston 2013), recent studies indicate that functional connectivity patterns are not stationary and fluctuate over very short periods of time on the order of seconds (Chang and Glover 2010, Handwerker, Roopchansingh et al. 2012, Hutchison, Womelsdorf et al. 2013, Parr, Rees et al. 2018). This has resulted in a new and rapidly evolving line of studies examining dynamic networks or time-varying functional connectivity (TVFC) patterns of the brain.

Studies of brain dynamics are critical for establishing a profound understanding of the brain given that the brain is a complex multiscale dynamic system rather than a stationary one (Lurie, Kessler et al. 2020). Dynamic brain networks have been associated with a wide range of cognitive and behavioral responses (Cole, Bassett et al. 2014, Shine, Bissett et al. 2016, Vidaurre, Hunt et al. 2018). More specifically, they have been used to determine the engagement of a subject in a specific cognitive task (Shirer, Ryali et al. 2012, Gonzalez-Castillo, Hoy et al. 2015), and have been associated with consciousness (Barttfeld, Uhrig et al. 2015, Godwin, Barry et al. 2015), learning (Bassett, Wymbs et al. 2011), and various neuropsychiatric and neurological disorders, such as schizophrenia (Sakoglu, Pearlson et al. 2010, Rashid, Arbabshirani et al. 2016), depression (Long, Cao et al. 2020, Martinez, Deco et al. 2020), Alzheimer’s disease (Jones, Vemuri et al. 2012, Gu, Lin et al. 2020), and Parkinson’s disease (Diez-Cirarda, Strafella et al. 2018, Zhu, Huang et al. 2019). New studies indicate that dynamic brain networks may provide more sensitive biomarkers for detecting differences between study populations or individuals than static networks (Rashid, Arbabshirani et al. 2016). In addition, studies of dynamic brain networks can answer more compelling open questions about cognitive and behavioral responses, as noted in (Lurie, Kessler et al. 2020).

Despite such insights, substantial challenges remain to enable more accurate analysis of dynamic brain networks and accurate interpretation of results. Development of multivariate statistical tools that allow for identifying associations between dynamic brain networks or TVFC patterns and phenotypic traits, as well as drawing inference from such associations, is among such critical challenges. Developing and disseminating explainable, validated multivariate statistical methods are paramount for relating phenotypic traits to dynamic changes in network properties of the brain, which will greatly aid in providing profound insights into normal and abnormal brain function. Dynamic changes in the systemic organization of brain networks confers much of our brains’ functional abilities (Buzsaki and Draguhn 2004, Bressler and Menon 2010). If functional connections are lost or rendered dynamically rigid due to an adverse health condition, compensatory connections may develop to maintain organizational consistency and functional abilities. Consequently, brain network analysis necessitates a suite of tools including a multivariate modeling framework for dynamic brain network data to assess effects of multiple variables of interest and topological network features on the overall network structure.

For the modeling framework, if we have

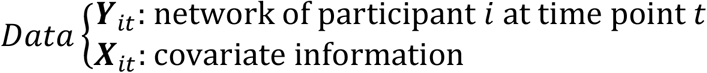

We wish to accurately estimate the probability density function of the networks given the covariates P(***Y***_*it*_|***X***_*it*_, ***β***_*it*_), where ***β***_*it*_ are the parameters that relate the covariates to the network structure as shown in Figure 1. However, the development of such methods has vastly lagged behind, mainly due to the same challenges which exist in developing multivariate statistical tools for static networks (Shehzad, Kelly et al. 2014, Simpson and Laurienti 2016, Bahrami, Laurienti et al. 2019). Most current methods rudimentarily compare the variability of connection strength or networks across study populations (Elton and Gao 2015, Fukushima, Betzel et al. 2018, Sizemore and Bassett 2018), failing to fully harness the wealth of information obtained via such a multivariate framework. As noted by (Shine, Breakspear et al. 2019), the neurobiological mechanisms underlying brain network dynamics (dynamic changes in functional architecture) remain poorly understood; and as pointed out in (Liu 2017) “novel methods are urgently needed for a better quantification of temporal dynamics in resting-state fMRI.” The development of rigorous statistical methods within a multivariate framework as described above are among such urgent needs.

**Figure 1.**
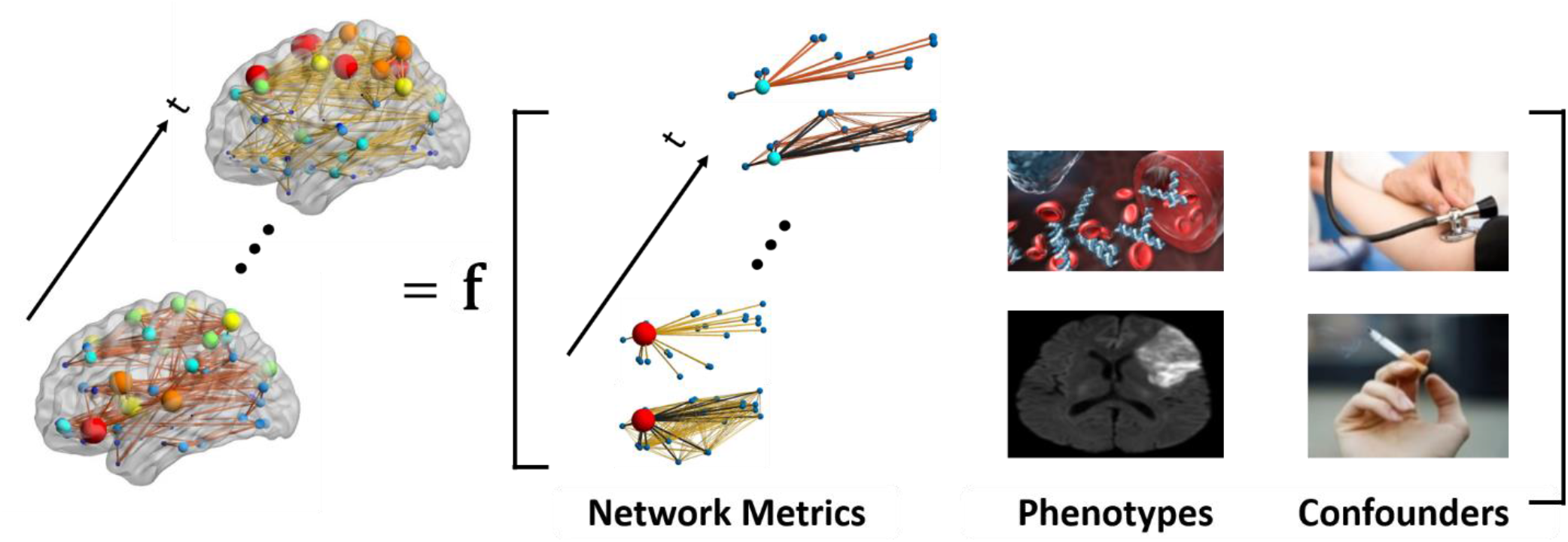
Dynamic brain networks as a function of endogenous and exogenous variables of interest. Dynamic patterns of brain connectivity (presence/absence and strength) is modeled as a function of (dynamic) nodal and global network variables (e.g., clustering coefficient, global efficiency, etc.) and exogenous covariates, including: phenotypes (e.g., blood measurements and brain damage) and possible confounding effects (e.g., hypertension and smoking).

Here we introduce a novel multivariate statistical framework for assessing phenotype-dynamic brain network pattern relationships and drawing inference from such relationships. This *model-based* framework is a fundamentally different approach toward analyzing dynamic brain networks when compared to current approaches which often use *data-driven* methodologies to identify “brain states” and their transitions with respect to health and behavioral outcomes (Allen, Damaraju et al. 2014, Vidaurre, Smith et al. 2017, Shine and Poldrack 2018, Shappell, Caffo et al. 2019). We develop this method by advancing a promising statistical mixed-modeling framework for static networks (Simpson and Laurienti 2015). Several extensions of the original framework (Bahrami, Laurienti et al. 2019, Simpson, Bahrami et al. 2019), as well as a Matlab toolbox with user-friendly graphical user interfaces (GUIs) (Bahrami, Laurienti et al. 2019) have recently been introduced. The original model and its extensions have been used in several studies (Bahrami, Laurienti et al. 2017, Burdette, Laurienti et al. 2020). However, it has yet to be extended to the dynamic network context. To our knowledge, this proposed extension will be the first to allow relating group- and individual-level characteristics to time-varying changes in spatial and topological brain network properties while also maintaining the capabilities of the original model, such as accounting for variance associated with confounders. We will demonstrate the utility of this framework in identifying the relationship between (fluid) intelligence and dynamic brain network patterns using 200 subjects from the HCP (Van Essen, Smith et al. 2013) study. Fluid intelligence (gF) refers to reasoning ability and the capacity of an individual to discern patterns or solve problems when that individual doesn’t have or has minimal resources or acquired knowledge to act upon (Cattell 1987). Understanding the neurobiological underpinnings of gF is of great interest, as it has been associated with a variety of cognitive abilities (Colom and Flores-Mendoza 2007, Unsworth, Fukuda et al. 2014, Ye, Li et al. 2019).

## 2. Materials and Methods

### 2.1 Motivating data

We used the rich data set provided by the HCP study (Van Essen, Smith et al. 2013) to be able to explore dynamic functional brain network differences in cognitively variable populations as a function of phenotype, while maintaining continuity with previous analyses to contrast and clearly distinguish the novel utilities of our proposed method. We specifically focused on demonstrating the utility of our framework in assessing the relationship between dynamic functional networks and intelligence due to the great interest in identifying such relationship. The HCP data released to date include 1200 individuals. Of those, 1113 (606 female; 283 minority) have complete MRI images, cognitive testing, and detailed demographic information. Participants in the HCP were screened to rule out neurological and psychiatric disorders. All data were collected on 3T Siemens MRI scanners located at Washington University or the University of Minnesota using identical scanning parameters. The HCP performed extensive testing and development to ensure comparable imaging at the two sites. The BOLD-weighted images were collected using the following parameters: TR = 720 ms, TE = 33.1 ms, voxel size 2 mm^3^, 72 slices, and 1200 volumes. In this study, we selected a subsample comprising 389 individuals with unique family identification numbers that also passed our image processing quality control assessments. For multiple individuals with the same family identification number, one individual was selected randomly. We initially used the entire 389 individuals, but we further reduced this to 200 individuals (randomly chosen from our subsample) after we faced convergence issues in modeling one of the two-part mixed-effect models. The HCP analyses are an exemplar; importantly, our methods can be applied to any network-based neuroimaging study.

### 2.2. Dynamic networks generation

We used minimally preprocessed rfMRI data from HCP (Glasser, Sotiropoulos et al. 2013). We used two scans for each individual, the left-to-right (LR) and right-to-left (RL). For each scan, we used ICA-AROMA (Pruim, Mennes et al. 2015) to correct for any motion artifact in the rfMRI data. A band-pass filter (0.009-0.08 Hz) was applied to each scan. The LR and RL scans for each individual were then concatenated temporally, and then a regression was performed with the mean tissue signals (GM, WM, and CSF), the six movement parameters and derivatives, as well as a binary regressor to account for any mean signal differences between the two groups (LR and RL scans). Our quality control process removed 116 individuals from the analysis. QC included manually checking the rfMRI for warping irregularities as well as remaining motion artifact after the above processing steps. For the remaining individuals, among those with the same family identification number, one individual was selected randomly. This provided a final dataset comprising 389 individuals with unique family identification numbers. For all 389 individuals, we divided the brain into 268 regions based on a functional atlas (Shen, Tokoglu et al. 2013), and averaged all time series within each region to create a single time-series for that region. We used a continuous wavelet transform (CWT) to filter artifact resulting from the LR and RL concatenation with a window size of 30s (covering 15s from the ending and starting points of LR and RL time series, respectively). We then prewhitened the time series to avoid undesired autocorrelation effects for our regression analyses and as recommended by (Honari, Choe et al. 2019) for dynamic network analyses using a sliding window correlation (SWC) approach.

Dynamic brain networks for each participant were constructed through a sliding window correlation approach. We used a modulated rectangular (mRect) window (Mokhtari, Akhlaghi et al. 2019) with a length of 120 volumes and the same shift size (i.e., 120 volumes) to generate non-overlapping windows. We understand that this is not a commonly used shift size as most studies use overlapping windows with 1 TR shift size; however, unlike other methods, we subsequently use the dynamic networks in a regression framework, thus we used non-overlapping windows to further reduce autocorrelation. The reasons and implications of our choices for window type, window length, and shift size will be further explained in the Discussion. The dynamic networks for each participant were generated by moving the window across the time series, and computing the Pearson’s correlation between time series of all pairs of 268 regions at each shift. This yielded 19 dynamic networks for each participant. We then thresholded all dynamic networks to remove negative correlations as multiple network measures, particularly clustering, remain poorly understood in networks with negative correlations (Telesford, Simpson et al. 2011, Friedman, Landsberg et al. 2014). Figure 2 shows a schematic exhibiting this dynamic network generation process.

**Figure 2.**
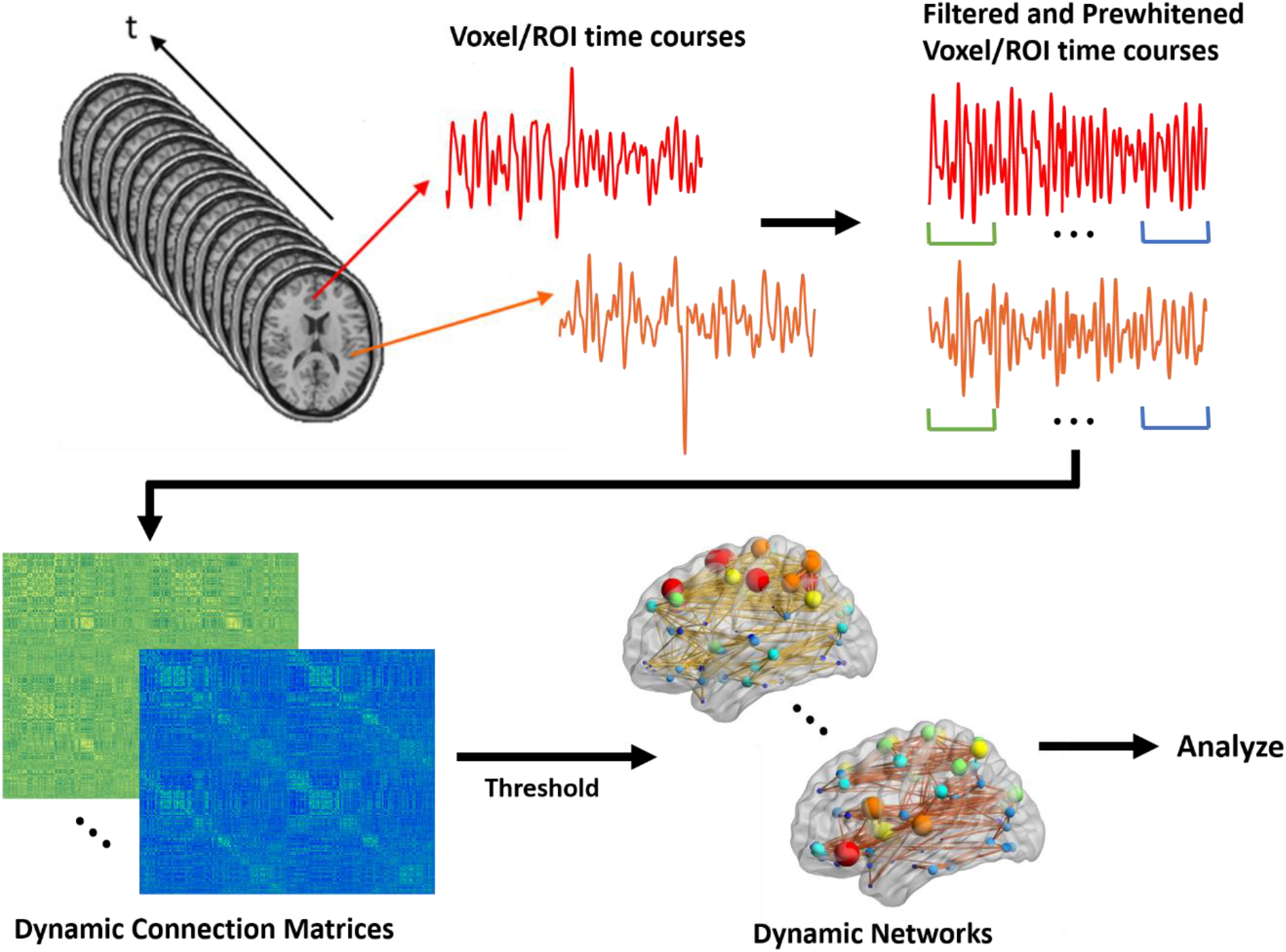
Schematic for generating dynamic brain networks from fMRI time series. Time series are first filtered and prewhitened to remove the undesired undershoots/overshoots in the middle as well as undesired effects of autocorrelation. Then, using a sliding window correlation (SWC) approach, functional connectivity between brain areas is estimated between all time series pairs at each shift to produce a connection matrix at that shift. By moving the window across the entire length of time series, a series of dynamic connection matrices will be produced for each participant. A threshold is applied to the matrix to remove negative connections. These networks are subsequently used for analyses.

We also used the original models (Simpson and Laurienti 2015) to conduct the same analyses but with static networks to see how the results are different with the results of the dynamic network analyses, and how the new introduced framework for dynamic networks provides more accurate results. For the static networks, we constructed a single network for each one of the 200 individuals by computing the correlation between average time series of all pairs of brain regions across the entire scanning time.

### 2.3. Mixed-effects modeling framework for weighted dynamic brain networks

Given that we have sparse weighted networks, a two-part mixed-effects model will be employed to model both the probability of a connection (presence/absence) and the strength of a connection, if it exists (Simpson and Laurienti 2015). The model includes the entire brain connectivity matrix of each participant, endogenous covariates, and exogenous covariates (see Figure 1). The endogenous covariates are summary variables extracted from the network to summarize global topology. The exogenous covariates are the biologically-relevant phenotypic variables (e.g. for our data, fluid intelligence, sex, race, and education among others). This statistical framework allows for the evaluation of group and individual effects. Another key feature of the model is the multivariate nature of the statistics. Inclusion of the actual connectivity matrices allows the statistics to be performed on the entire network simultaneously, rather than performing edge-by-edge analyses in a massively univariate fashion.

More specifically, let *Y*_*ijkt*_represent the *strength* of the connection (quantified as the correlation in our case) and *R*_*ijkt*_indicate whether a connection is present (*presence* variable) between node *j* and node *k* for the *i*^*th*^ subject at time *t*. Thus, *R*_*ijkt*_ = 0 if *Y*_*ijkt*_ = 0, and *R*_*ijkt*_ = 1 if *Y*_*ijkt*_ > 0 with conditional probabilities

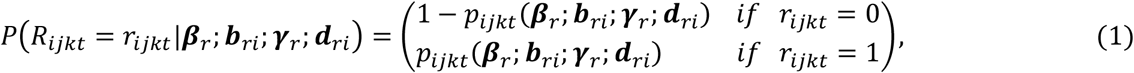

where *p*_*ijkt*_ (***β***_*r*_; ***b***_*ri*_; ***γ***_*r*_) is the probability of a connection between nodes *j* and *k* for subject *i* at time *t*. We then have the following logistic mixed model (part I model) for the probability of this connection:

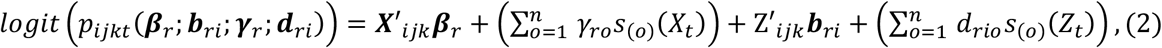

where ***β***_*r*_ is a vector of population parameters (fixed effects) that relate the probability of a connection to a set of covariates (***X***_*ijk*_) for each subject and nodal pair (dyad), ***b***_*ri*_ is a vector of subject- and node-specific parameters (random effects) that capture how this relationship varies about the population average (***β***_*r*_) by subject and node (***Z***_*ijk*_), 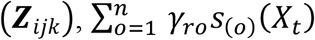 corresponds to a population-level *n*^*th*^ order orthonormal polynomial model capturing the dynamic trend in the *presence* of connections across time, and 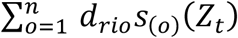 corresponds to an individual-level *N*^th^ order orthonormal polynomial model capturing how much the subject-specific trends deviate from the population trend. Employing an orthonormal polynomial model in this manner has been shown to accurately represent the trend in time series data while avoiding the computational issues resulting from the use of natural polynomials (Simpson and Edwards 2013, Edwards and Simpson 2014).

For the part II model, which aims to model the strength of a connection given that there is one, we let *S*_*ijkt*_ = [*Y*_*ijkt*_ |*R*_*ijkt*_ = 1]. In our case, the *S*_*ijkt*_ will be the values of the correlation coefficients between nodes *j* and *k* for subject *i* at time *t*. We can then use Fisher’s Z-transform, denoted as *FZT*, to induce normality for the following mixed model (part II model)

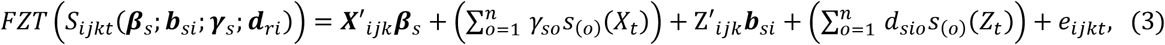

where ***β***_*s*_ is a vector of population parameters that relate the strength of a connection to the same set of covariates (***X***_*ijk*_) for each subject and nodal pair (dyad), ***b***_*si*_ is a vector of subject- and node-specific parameters that capture how this relationship varies about the population average (***β***_*s*_) by subject and node (***Z***_*ijk*_), 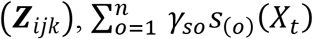 corresponds to a population-level *n*^*th*^ order orthonormal polynomial model capturing the dynamic trend in the *strength* of connections across time, 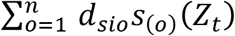 corresponds to an individual-level *n*^*th*^ order orthonormal polynomial model capturing how much the subject-specific trends deviate from the population trend, and *e*_*ijkt*_ accounts for the random noise in the connection strength of nodes *j* and *k* for subject *i* at time *t*.

In this study, the covariates (***X***_*ijk*_) used to explain and predict both the presence and strength of connection are: 1) *Net*: the average of the following network variables (categorized and further detailed in Table 1 below and in (Rubinov and Sporns 2010, Simpson, Bowman et al. 2013) in each dyad: Clustering Coeficient (), Global Efficiency (*g ob*), Degree (*k*) (difference between connected nodes instead of average to capture “assortativity”), Modularity (*Q*), and Leverage Centrality (*l*); 2) *COI*: Covariates of Interest (fluid intelligence (gF) in our study – we modeled gF as a continuous covariate. gF in the HCP protocol has been assessed using the Raven’s progressive matrices with 24 items with scores being integers representing number of correct items (Bilker, Hansen et al. 2012)); 3) *Int*: Interactions of the Covariate of Interest with the variables in 1); and 4) *Con*: Confounders (for our data, Sex (binary), Age (continuous), years of Education (categorical with three levels – level 1 (≤11), level 2 (12-16), and level 3 (≥17)), BMI (continuous), Race (categorical with six categories – cat 1 (Am. Indian/Alaskan Nat.), cat 2 (Asian/Nat. Hawaiian/Other Pacific Is.), cat 3 (Black or African Am.), cat 4 (White), cat 5 (More than one), cat 6 (Unknown or Not Reported)), Ethnicity (categorical with three categories – cat 1 (Hispanic/Latino), cat 2 (Not Hispanic/Latino), cat 3 (Unknown or Not Reported)), Handedness (continuous – ranging from −100 to +100, with negative and positive numbers indicating whether participants were more left- or right-handed, respectively assessed using the Edinburgh Handedness Inventory (EHI) (Oldfield 1971), Income (Continuous – Toal household income), Alcohol abuse (Binary – Indicating whether participant met DSM4 criteria for alcohol abuse), Alcohol dependence (Binary – Indicating wether participant met DSM4 criteria for alcohol dependence), Smoking status (Binary – Indicating whether pariticipant smoked or not), Spatial distance between nodes (importance of spatial distance as potential geometric confounders has been discussed in (Friedman, Landsberg et al. 2014)), and square of spatial distance between nodes). Thus, we can decompose ***β***_*r*_ and ***β***_*s*_ into ***β***_*r*_ = [***β***_*r*,0_***β***_*r,net*_ ***β***_*r,coi*_***β***_*r,int*_ ***β***_*r,con*_] and ***β***_*s*_ = [***β***_*s*,0_ ***β***_*s,net*_ ***β***_*s,coi*_ ***β***_*s,int*_ ***β***_*s,con*_] to correspond with the population intercepts and these covariates. For the random-effects vectors we have that ***b***_*ri*_ = [*b*_*ri*,0_ ***b***_*ri,net*_ ***b***_*ri,dist*_ ***δ***_*ri,j*_***δ***_*ri,k*_] and ***b***_*si*_ = [*b*_*si*,0_ ***b***_*si,net*_ ***b***_*si,dist*_ ***δ***_*si,j*_***δ***_*si,k*_], where *b*_*ri*,0_ and *b*_*si*,0_ quantify the deviation of subject-specific intercepts from the population intercepts (***β*** _*r,0*_ and ***β*** _*s,0*_), ***b***_*ri,net*_ and ***b***_*si,net*_ contain the subject-specific parameters that capture how much the relationships between the network variables in 1) and the *presence* and *strength* of a connection vary about the population relationships (***β*** _*r,net*_ and ***β*** _*s,net*_), respectively, ***b***_*ri,dist*_ and ***b***_*si,dist*_ contain the subject-specific parameters that capture how much the relationship between spatial distance (and square of spatial distance) and the *presence* and *strength* of a connection vary about the population relationships respectively, ***δ***_*ri,j*_ and ***δ***_*si,j*_ contain nodal-specific parameters that represent the propensity for node *j* (of the given dyad) to be connected and the magnitude of its connections, respectively, and ***δ***_*ri,k*_ and ***δ***_*si,k*_ contain nodal-specific parameters that represent the propensity for node *k* (of the given dyad) to be connected and the magnitude of its connections respectively. **Parameters for all *T* time points (number of networks per individual)** (***t*** = **1, 2,…**, ***T***) **are estimated or predicted simultaneously from the model**. In general, additional covariates can also be incorporated as guided by the biological context.

**Table 1.** Network measures by category

| Category | Measure(s) |
| --- | --- |
| Functional segregation | Clustering coefficient |
| Functional integration | Global efficiency |
| Resilience | Degree difference |
| Centrality and information flow | Leverage centrality |
| Community structure | Modularity |

Specifying a reasonable covariance model (balancing appropriate complexity with parsimony and computational feasibility) is paramount for a unified dynamic multitask model such as the one developed here. Toward this end, we assume that ***b***_*ri*_, ***d***_*ri*_, ***b***_*si*_, ***d***_*si*_, and ***e***_*i*_ are normally distributed and mutually independent, with variance component covariance structures for ***b***_*ri*_, ***d***_*ri*_, ***b***_*si*_, and ***d***_*si*_, and the standard conditional independence structure for ***e***_*i*_. That is, ***b***_*ri*_ *∼ N*(***0, Σ***_*bri*_(***τ***_*br*_) = *diag*(***τ***_*br*_)), where 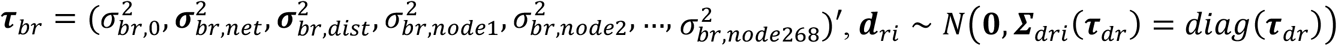 where 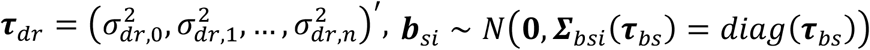 where 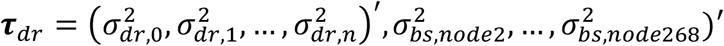 and ***d***_*si*_ *∼ N*(***0, Σ***_*dsi*_ (***τ***_*ds*_) = *diag*(***τ***_*ds*_)) where 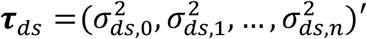 —yielding (276 + (*n* + 1)) random effects variance parameters for both the *presence* and *strength* models—and ***e***_*i*_ *∼ N*(***0, Σ***_*ei*_ = ***σ***^2^***I***). These variance and covariance parameters will provide insight into whether individual and group differences in variability in dynamics relate to health and behavioral outcomes. Parameter estimation is conducted via restricted pseudo-likelihood (Wolfinger and Oconnell 1993) with the residual approximation of the *F*-test for a Wald statistic employed for inference.

We implemented the models (eqs. 2 and 3) above to describe and compare brain network dynamics as a function of fluid intelligence. For both models, we started model fitting with the entire set of random-effects, i.e., random effects for: intercept, nodal network measures (clustering, global efficiency, degree, and leverage centrality), distance, and nodal propensities. However, after facing convergence issues, we dropped nodal propensities from our random effects. We assessed model goodness-of-fit (GOF) and consistency of estimates (to further avoid overfitting) to determine the orthonormal polynomial degree yielding the best model fits. We fit the two-part model defined above with the mentioned fixed- and random-effect parameters using orthonormal polynomial models of degrees ranging from 3-18 (giving 16 model fits), and determined the “best” model based on a composite approach employing the Akaike Information Criterion (AIC) (Akaike 1981), Bayesian Information Criterion (BIC) (Schwarz 1978), modified AIC (AICc) (Hurvich and Tsai 1989), Hannan-Quinn Information Criterion (HQIC) (Hannan and Quinn 1979), and Consistent AIC (CAIC) (Bozdogan 1987) GOF measures as well as the consistency of the obtained parameter estimates and p-values to further avoid overfitting. We used Matlab to generate the appropriate data frame for our framework and used SAS v9.4 on a Linux operating system with 330 GB of RAM and 2.60 GH processor to perform the model fitting.

## 3. Results

Here, we show our framework’s ability in identifying the relationship between fluid intelligence and dynamic brain networks. For orthonormal polynomial models of degrees ranging from 3-18, all GOF measures for the strength model (eq.3) slightly improved with increasing degree, providing good fits for almost all degrees. However, the models with polynomial degrees ranging from 9-16 provided the most consistent estimates and p-values. Thus, to avoid overiftting while still using a model with a relatively good fit as indicated by the GOF measures, we used the model with polynomial degree of 12 as a middle ground between 9-16. For the probability model, although all GOF measures slightly improved with increasing degree too, the differences were negligible. Thus, we used the same polynomial degree (12) for consistency. The estimates, standard errors, and p-values for the polynomial parameters are presented in Table 2.

**Table 2.** Fixed-effect estimates, SEs, and P-values for 12^th^ degree orthornormal polynomial fit

| Models | Parameters | Ortho Poly degree | Estimate | SE | P-value |
| --- | --- | --- | --- | --- | --- |
| **Probability Model** | $\beta_{r,0}$ | Intercept | -0.0325 | 0.0029 | <0.0001 |
| | $\beta_{r,1}$ | 1 | -12.864 | 13.098 | 0.3260 |
| | $\beta_{r,2}$ | 2 | 3.2334 | 11.666 | 0.7817 |
| | $\beta_{r,3}$ | 3 | -45.319 | 13.146 | 0.0006 |
| | $\beta_{r,4}$ | 4 | 64.035 | 11.513 | <0.0001 |
| | $\beta_{r,5}$ | 5 | -15.858 | 12.414 | 0.2015 |
| | $\beta_{r,6}$ | 6 | -13.590 | 11.848 | 0.2514 |
| | $\beta_{r,7}$ | 7 | -31.703 | 11.722 | 0.0068 |
| | $\beta_{r,8}$ | 8 | 21.245 | 10.751 | 0.0482 |
| | $\beta_{r,9}$ | 9 | 10.496 | 10.983 | 0.3393 |
| | $\beta_{r,10}$ | 10 | -11.572 | 11.885 | 0.3302 |
| | $\beta_{r,11}$ | 11 | -28.431 | 11.331 | 0.0121 |
| | $\beta_{r,12}$ | 12 | 20.729 | 11.242 | 0.0652 |
| **Strength Model** | $\beta_{s,0}$ | Intercept | 0.3297 | 0.0011 | <0.0001 |
| | $\beta_{s,1}$ | 1 | -17.061 | 4.779 | 0.0004 |
| | $\beta_{s,2}$ | 2 | 2.7768 | 3.977 | 0.4851 |
| | $\beta_{s,3}$ | 3 | -26.615 | 4.228 | <0.0001 |
| | $\beta_{s,4}$ | 4 | 33.522 | 3.892 | <0.0001 |
| | $\beta_{s,5}$ | 5 | -2.0592 | 3.677 | 0.5755 |
| | $\beta_{s,6}$ | 6 | -1.0681 | 4.021 | 0.7905 |
| | $\beta_{s,7}$ | 7 | -19.033 | 3.949 | <0.0001 |
| | $\beta_{s,8}$ | 8 | 20.267 | 4.156 | <0.0001 |
| | $\beta_{s,9}$ | 9 | 7.2895 | 4.244 | 0.0858 |
| | $\beta_{s,10}$ | 10 | -8.2756 | 4.051 | 0.0411 |
| | $\beta_{s,11}$ | 11 | -13.659 | 3.774 | 0.0003 |
| | $\beta_{s,12}$ | 12 | 2.5260 | 3.665 | 0.4907 |

The parameter estimates, standard errors, and p-values (based on the residual approximation of the F-test for a Wald statistic) associated with the fixed-effect covariates are presented in Table 3. The estimates quantify the relationship between dynamic patterns of probability (presence/absence) and strength of (present) connections between nodes (brain regions), as dependent variables, and the previously mentioned sets of covariates, including (dynamic patterns of) endogenous network measures, fluid intelligence as our covariate of interest, and confounders (sex, age, education, BMI, race, ethnicity, handedness, income, DSM4 alcohol abuse, DSM4 alcohol dependence, smoking status, spatial distance and square of spatial distance between nodes). The estimates for interaction covariates shows if (and how) the relationship between dynamic patterns of probability/strength of connections and dynamic patterns of endogenous network measures are affected by fluid intelligence. Notable results are detailed in the following sections.

**Table 3.**
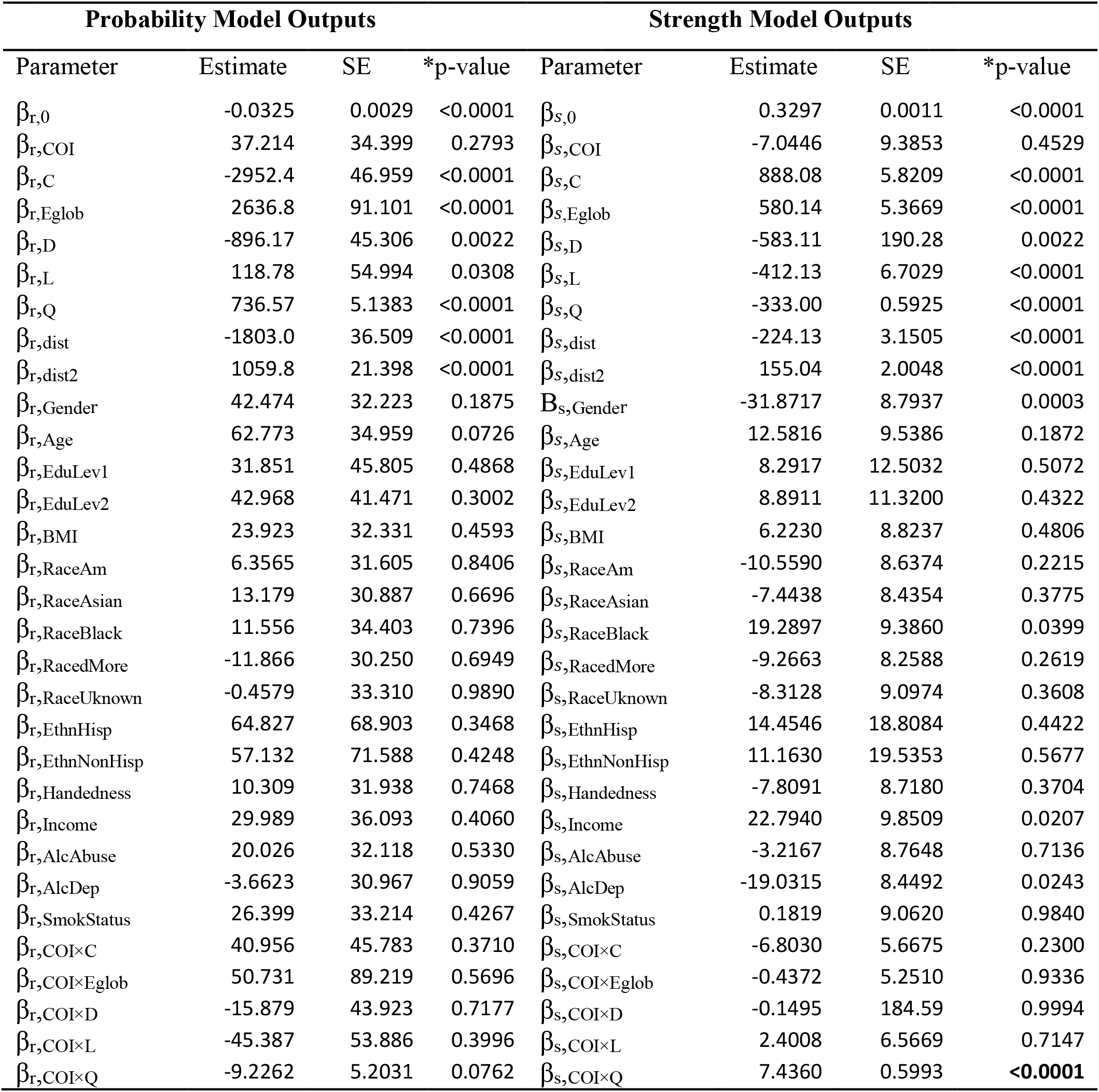
Parameter estimates, standard errors, and p-values for dynamic networks

### 3.1 Dynamic network analysis

#### 3.1.1. Endogenous network measures and confounding covariates

As Table 3 presents, dynamic changes of clustering (functional segregation), global efficiency (functional integration), degree difference (functional resilience), and leverage centrality (information flow), all play important roles in explaining dynamic patterns of both connection probability and strength. Among the confounding covariates, spatial distance and square of spatial distance are important covariates in explaining the dynamic patterns of both connection probability and strength, while gender, race (Black or American-African as compared with white (reference group)), income, and alcohol dependence are important in explaining the dynamic patterns of just connection strength.

#### 3.1.2. Fluid intelligence

Fluid intelligence, our covariate of interest (COI), is neither directly related to dynamic patterns of connection probability (presence/absence) nor connection strength as indicated by the p-values associated with ***β***_*r,COI*_ (p-value = 0.2793) and ***β***_*s,COI*_ (p-value = 0.4529), respectively. However, it has a significant effect on the relationship between dynamic changes of connection strength and dynamic changes of whole-brain modularity as indicated by the p-value associated with ***β***_*s,COI*×*Q*_ (p-value <0.0001), and a marginally significant effect on the relationship between dynamic changes of connection probability and dynamic changes of whole-brain modularity as indicated by the p-values associated with ***β***_*r,COI*×*Q*_ (p-value = 0.0762), while having no effect on other relationships. Dynamic changes of whole-brain modularity and connection strength are negatively associated with each other (***β***_*s,Q*_), which implies the dominance of between-community (rather than within-community) connections in driving the dynamic changes of whole-brain modularity. Fluid intelligence interacts with this relationship — as intelligence increases, dynamic changes of modularity are less driven by dynamic changes in between-community (and more by within-community) connectivity as indicated by the positive and significant estimate for ***β***_*s,COI*×*Q*_. (However, the dynamics of between-community connections are still the dominant factors in driving the dynamic changes of whole-brain modularity.) Our results might imply that brain networks in people with higher fluid intelligence are more flexible with respect to changes in their modularity at rest. These changes are associated with both stronger within-community connections (more specialized neural communities) and weaker between-community connections (more segregated neural communities). This is illustrated in Figure 3.

**Figure 3.**
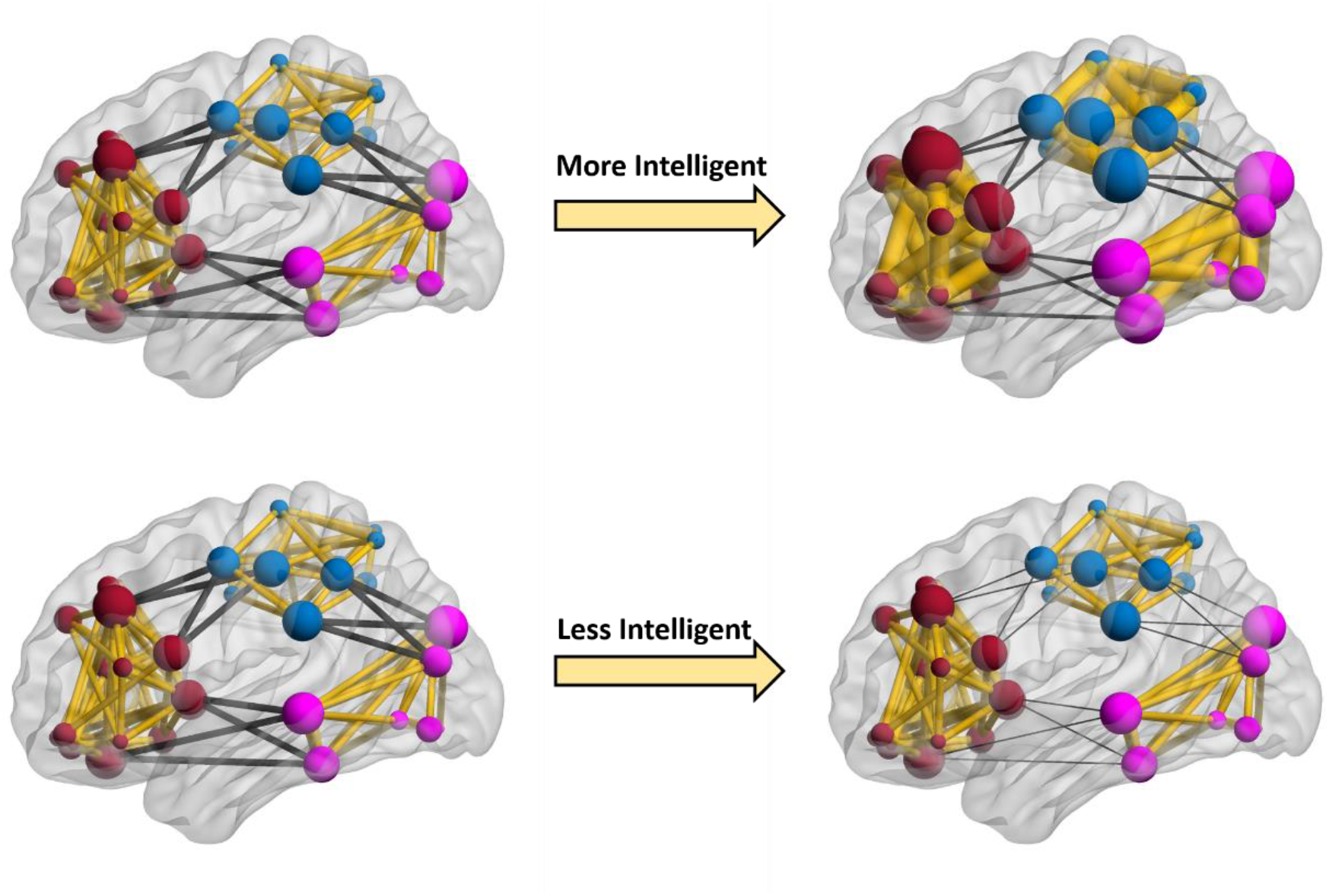
Cartoon depiction of fluid intelligence effects on dynamic brain networks. The nodes represent brain regions, and edges represent dynamic functional connections. To illustrate the effects of fluid intelligence on dynamic changes of modularity as interpreted from Table 3, three random communities marked with separate colors, dark red, light blue, and purple, are shown in this figure. The within- and between-community connections are shown with the yellow and black colors, respectively. As this figure illustrates, dynamic changes of modularity is predominantly determined by between-community connections for any level of intelligence (here two level is shown – low and high). However, when comparing the more and less intelligent participants (networks on the right), in more intelligent participants, dynamic changes of modularity is less determined by between-community connections (thicker dark edges in top right), and dynamic changes of within-community connections also play more important roles in changing the modularity (thicker yellow edges in top right network). We should note that fluid intelligence was modeled as a continuous variables, and here, only for simplicity, we have shown how intelligence affects modularity for two extreme conditions.

### 3.2. Dynamic versus Static network analysis

To further demonstrate how the developed framework for dynamic brain networks provides a better fit and different insight into effects of phenotypic traits on brain networks, we conducted an additional analysis to examine the effects of fluid intelligence on static brain networks for comparison. We used the original mixed-effects models introduced in (Simpson and Laurienti 2015) with the exact same fixed- and random-effects, and the same random-effects variance covariance structure as those used for dynamic networks. All GOF measures indicate much better fits for dynamic brain networks (Tables 4, 5), probably due to accounting for the dynamic trends in the brain network data. Also, the results for modeling the static brain networks presented in Table 6 clearly illustrate that different conclusions would be drawn from employing this less favorable modeling approach, particularly when comparing the effects of fluid intelligence on brain modularity and connectivity. While dynamic patterns of whole-brain modularity are modified by fluid intelligence, the static network analyses indicate no such effect.

**Table 4.** Probability model’s GOF measures for dynamic and static network analyses

| Analysis | -2 Res Log Likelihood | Generalized Chi-Square |
| --- | --- | --- |
| Dynamic (ortho poly degree 12) | 5.76E8 | 1.36E8 |
| Static | 0.31E8 | 0.07E8 |

**Table 5.** Strength model’s GOF measures for dynamic and static network analyses

| Analysis | AIC | AICC | BIC | CAIC | HQIC |
| --- | --- | --- | --- | --- | --- |
| Dynamic (ortho poly degree 12) | -7448537 | -7448537 | -7448471 | -7448451 | -7448510 |
| Static | -3640945 | -3640945 | -3640918 | -3640910 | -3640934 |

**Table 6.** Parameter estimates, standard errors, and p-values for static networks

| Probability Model Outputs |  |  |  | Strength Model Outputs |  |  |  |
| --- | --- | --- | --- | --- | --- | --- | --- |
| Parameter | Estimate | SE | *p-value | Parameter | Estimate | SE | *p-value |
| $\beta_{r,0}$ | -0.2584 | 0.0809 | 0.0014 | $\beta_{s,0}$ | 0.1022 | 0.0289 | 0.0004 |
| $\beta_{r,COI}$ | -0.2073 | 0.0110 | 0.2793 | $\beta_{s,COI}$ | -0.0089 | 0.0043 | 0.0369 |
| $\beta_{r,C}$ | -2952.4 | 46.959 | <0.0001 | $\beta_{s,C}$ | 0.0777 | 0.0008 | <0.0001 |
| $\beta_{r,Eglob}$ | -0.0403 | 0.0079 | <0.0001 | $\beta_{s,Eglob}$ | 0.0700 | 0.0009 | <0.0001 |
| $\beta_{r,D}$ | 0.0176 | 0.0042 | <0.0001 | $\beta_{s,D}$ | -0.0433 | 0.0005 | <0.0001 |
| $\beta_{r,L}$ | 0.1714 | 0.0092 | <0.0001 | $\beta_{s,L}$ | -0.0419 | 0.0009 | <0.0001 |
| $\beta_{r,Q}$ | 0.0918 | 0.0110 | <0.0001 | $\beta_{s,Q}$ | -0.0177 | 0.0039 | <0.0001 |
| $\beta_{r,dist}$ | -0.2993 | 0.0050 | <0.0001 | $\beta_{s,dist}$ | -0.0398 | 0.0005 | <0.0001 |
| $\beta_{r,dist2}$ | 0.1510 | 0.0024 | <0.0001 | $\beta_{s,dist2}$ | 0.0211 | 0.0003 | <0.0001 |
| $\beta_{r,Gender}$ | 0.0022 | 0.0220 | 0.9197 | $\beta_{s,Gender}$ | 0.0121 | 0.0079 | 0.1263 |
| $\beta_{r,Age}$ | 0.0025 | 0.0119 | 0.8350 | $\beta_{s,Age}$ | 0.0043 | 0.0043 | 0.3092 |
| $\beta_{r,EduLev1}$ | 0.0524 | 0.0415 | 0.2070 | $\beta_{s,EduLev1}$ | 0.0018 | 0.0149 | 0.9062 |
| $\beta_{r,EduLev2}$ | 0.0164 | 0.0298 | 0.5822 | $\beta_{s,EduLev2}$ | 0.0101 | 0.0107 | 0.3471 |
| $\beta_{r,BMI}$ | 0.0232 | 0.0114 | 0.0411 | $\beta_{s,BMI}$ | -0.0086 | 0.0041 | 0.0348 |
| $\beta_{r,RaceAm}$ | 0.0967 | 0.1523 | 0.5253 | $\beta_{s,RaceAm}$ | -0.0816 | 0.0549 | 0.1374 |
| $\beta_{r,RaceAsian}$ | -0.0038 | 0.0400 | 0.9250 | $\beta_{s,RaceAsian}$ | -0.0107 | 0.0144 | 0.4570 |
| $\beta_{r,RaceBlack}$ | -0.0317 | 0.0359 | 0.3777 | $\beta_{s,RaceBlack}$ | 0.0157 | 0.0129 | 0.2234 |
| $\beta_{r,RacedMore}$ | 0.1128 | 0.0842 | 0.1803 | $\beta_{s,RacedMore}$ | -0.0109 | 0.0304 | 0.7209 |
| $\beta_{r,RaceUnknown}$ | 0.0248 | 0.0817 | 0.7618 | $\beta_{s,RaceUnknown}$ | 0.0043 | 0.0295 | 0.8843 |
| $\beta_{r,EthnHisp}$ | -0.0965 | 0.0808 | 0.2320 | $\beta_{s,EthnHisp}$ | 0.0464 | 0.0289 | 0.1083 |
| $\beta_{r,EthnNonHisp}$ | -0.0580 | 0.0769 | 0.4507 | $\beta_{s,EthnNonHisp}$ | 0.0393 | 0.0275 | 0.1536 |
| $\beta_{r,Handedness}$ | 0.0081 | 0.0109 | 0.4531 | $\beta_{s,Handedness}$ | -0.0041 | 0.0039 | 0.2915 |
| $\beta_{r,Income}$ | -0.0098 | 0.0122 | 0.4266 | $\beta_{s,Income}$ | 0.0042 | 0.0044 | 0.3429 |
| $\beta_{r,AlcAbuse}$ | 0.0311 | 0.0279 | 0.2637 | $\beta_{s,AlcAbuse}$ | 0.0063 | 0.0100 | 0.5313 |
| $\beta_{r,AlcDep}$ | -0.0162 | 0.0402 | 0.6870 | $\beta_{s,AlcDep}$ | -0.0082 | 0.0145 | 0.5726 |
| $\beta_{r,SmokStatus}$ | -0.0628 | 0.0307 | 0.0409 | $\beta_{s,SmokStatus}$ | 0.0034 | 0.0110 | 0.7616 |
| $\beta_{r,COI \times C}$ | -0.0069 | 0.0111 | 0.5333 | $\beta_{s,COI \times C}$ | 0.0009 | 0.0008 | 0.2866 |
| $\beta_{r,COI \times Eglob}$ | 0.0026 | 0.0079 | 0.7475 | $\beta_{s,COI \times Eglob}$ | -0.0009 | 0.0009 | 0.2927 |
| $\beta_{r,COI \times D}$ | -0.0016 | 0.0042 | 0.7117 | $\beta_{s,COI \times D}$ | -0.0005 | 0.0005 | 0.2367 |
| $\beta_{r,COI \times L}$ | 0.0033 | 0.0092 | 0.7226 | $\beta_{s,COI \times L}$ | -0.0012 | 0.0009 | 0.1988 |
| $\beta_{r,COI \times Q}$ | 0.0055 | 0.0117 | 0.6401 | $\beta_{s,COI \times Q}$ | -0.0024 | 0.0042 | 0.5592 |

## 4. Discussion

As the interest in dynamic brain networks continues to grow, new methods are needed to enable gleaning neurobiological insight into this complex and big data. Development of multivariate statistical methods, particularly model-based ones, which allow quantifying relationships between phenotypic traits and dynamic patterns of brain connectivity and topology and drawing inference from such relationships is among the urgent needs. Development of such methods even for static networks has remained a challenge given the size, complexity, and multiscale dependence inherent in brain network data. However several model-based methods (Simpson, Hayasaka et al. 2011, Shehzad, Kelly et al. 2014, Simpson and Laurienti 2015) and various data-driven multivariate methods (Calhoun, Adali et al. 2001, Beckmann and Smith 2004, Allen, Erhardt et al. 2011, Smith, Hyvarinen et al. 2014) have been introduced and extensively used for static networks. Dynamic changes in the systematic organization of our brain networks confer much of our brains’ functions abilities due to the fact that our brain is a complex multiscale dynamic system with known and unknown compensatory mechanisms at multiple scales. Thus, methods that allow analyzing the brain within a multivariate framework can provide much deeper insights into dynamic patterns of brain networks in health and disease. In addition, multivariate model-based tools enable aligning neuroscientific hypotheses with the analytic approach which is ideal for dynamic brain network analysis (Preti, Bolton et al. 2017). Nevertheless, no model-based multivariate method has been introduced for dynamic network analyses to our knowledge.

Here we provided a baseline model-based multivariate method to relate phenotypic traits to dynamic patterns of brain connectivity and topology. We developed this model by advancing a two-part mixed-effects regression framework for static brain networks (Simpson and Laurienti 2015). Our proposed model allows accounting for the connectivity/network dynamics when assessing group differences and phenotype-health outcome relationships, to avoid confounding and drawing erroneous conclusions. Most current methods used to assess dynamic brain networks reduce this data into dynamic patterns of individual brain connections (Simony, Honey et al. 2016, Schmlazle, O’Donnell et al. 2017, Tewarie, Liuzzi et al. 2019) or commonly used summary network variables, such as node degree or modularity (Jones, Vemuri et al. 2013, Kabbara, Khalil et al. 2019) rather than analyzing the systemic dynamics of the brain networks. Such methods not only fail to model the brain as a multiscale dynamic system (Lurie, Kessler et al. 2020), but often entail matching study populations to perform group comparisons, which is a daunting task for most neuroimaging studies. Our model provides a framework to assess the systemic dynamics of brain networks and thus to account for complex dynamics of the brain via the simultaneous modeling of brain connectivity and topological network variables. The multivariate nature of this framework reduces demands for matching study populations as any number of confounding effects can be incorporated as covariates, and the effects of multiple covariates of interest can be studied in a single model.

We demonstrated the utility of our model in identifying the relationship between fluid intelligence and dynamic patterns of brain connectivity and topological network variables using the rich data set provided by the HCP study (Van Essen, Smith et al. 2013). Our model allowed accounting for various sources of potential confounding effects, such as sex, education, age, and alcohol abuse, among others. Our results indicated that dynamic patterns of brain modularity and connection strength are significantly affected by fluid intelligence. More specifically, our results showed that for any level of fluid intelligence, dynamic patterns of modularity are predominantly associated with between-community, rather than within-community, connections. However, fluid intelligence modulates this trend such that, across an entire spectrum of fluid intelligence, dynamics of whole-brain modularity play a less important role in driving changes in between-community connections for higher fluid intelligence values (with dynamics of within-community connections probably being affected more). While the ultimate neurobiological interpretations of such effects is speculative at this point, our results may suggest that brain networks in more intelligent participants are more flexible with respect to changes in its modularity at rest, such that dynamic patterns of modularity are determined by both establishing stronger within-community connections (more specialized neural communications) and weaker between-community connections (more segregated neural communities). Other studies have reported associations between brain modularity and intelligence (Chaddock-Heyman, Weng et al. 2020), as well as significant correlations between creativity and learning and dynamic patterns of brain modularity (Bassett, Wymbs et al. 2011, Kenett, Betzel et al. 2020).

We used a sliding window correlation approach in this analysis as it has remained the most popular approach to examine dynamic brain networks (Hutchison, Womelsdorf et al. 2013, Allen, Damaraju et al. 2014, Rashid, Damaraju et al. 2014, Preti, Bolton et al. 2017, Bahrami, Lyday et al. 2019). However, in the absence of a “gold-standard” the optimal choice for window type, window length, and step size is challenging. We used a modulated rectangular (mRect) window due to its superior performance in examining dynamic brain networks when compared to other conventional window types (Mokhtari, Akhlaghi et al. 2019). The window lengths used in the literature commonly range from the 30s to 240s (Chang and Glover 2010, Kiviniemi, Vire et al. 2011, Handwerker, Roopchansingh et al. 2012, Kucyi and Davis 2014, Mokhtari, Rejeski et al. 2018). We used 120 volumes for our window length for multiple reasons, including: i) to provide more stabilized correlation values while not losing the variability of the brain dynamics, ii) due to its wider use which makes comparing and contrasting our method with currently used methods easier, and iii) model fit and convergence considerations of our proposed method. It is also important to note that no commonly used window length can accurately identify different states of correlation (Shakil, Lee et al. 2016), and that the SWC is only used to demonstrate the utility of our method rather than to provide comprehensive analyses of fluid intelligence-dynamic brain network associations. Also, typical shift sizes used in the literature range from 1 TR to 50% of the window length (Chang, Liu et al. 2013, Kucyi and Davis 2014, Shakil, Magnuson et al. 2014), with the 1 TR being the most commonly used shift size (Shakil, Lee et al. 2016). However, as our proposed method subsequently uses the dynamic networks in a regression framework, we used a shift size of 120 volumes, equal to the window length, to create non-overlapping windows and thus further reduce autocorrelation. Additionally, using non-overlapping windows allowed using a smaller, but sufficient number of dynamic networks for each participant and thus helped avoiding possible convergence issues.

Our work here is not without limitations. The proposed mixed-effects framework can be used for predicting dynamic networks based on participant characteristics as well as simulating dynamic networks from the modeled distributions and desired characteristics. However, we have not demonstrated these capabilities here. These capabilities will be demonstrated in future work given that they require extensive analytical assessment and exposition which lie beyond the scope of this paper. We will also make our proposed framework accessible to neuroimaging researchers by incorporating new GUIs into WFU_MMNET, the software developed for the application of the original static model (Bahrami, Laurienti et al. 2019). Future studies will also assess the effects of the sliding window parameters (e.g., window length and shift size) as well as the parcellation choice on model results (Power, Cohen et al. 2011, Bahramf and Hossein-Zadeh 2014, Glasser, Coalson et al. 2016).

## Acknowledgements

This work was supported by National Institute of Biomedical Imaging and Bioengineering (R01EB024559) and National Institute of Environmental Health Sciences (R01 ES00873922S1).

